# Modelling haplotypes with respect to reference cohort variation graphs

**DOI:** 10.1101/101659

**Authors:** Yohei Rosen, Jordan Eizenga, Benedict Paten

## Abstract

Current statistical models of haplotypes are limited to panels of haplotypes whose genetic variation can be represented by arrays of values at linearly ordered bi- or multiallelic loci. These methods cannot model structural variants or variants that nest or overlap. A variation graph is a mathematical structure that can encode arbitrarily complex genetic variation. We present the first haplotype model that operates on a variation graph-embedded population reference cohort. We describe an algorithm to calculate the likelihood that a haplotype arose from this cohort through recombinations and demonstrate time complexity linear in haplotype length and sublinear in population size. We furthermore demonstrate a method of rapidly calculating likelihoods for related haplotypes. We describe mathematical extensions to allow modelling of mutations. This work is an essential step forward for clinical genomics and genetic epidemiology since it is the first haplotype model which can represent all sorts of variation in the population.

## 1 Background

Statistical modelling of individual haplotypes within population distributions of genetic variation dates back to Kingman’s *n-coalescent* [5]. In general, the coalescent and other models describe haplotypes as generated from some structured state space via recombination and mutation events.

While coalescent models are powerful generative tools, their computational complexity is unsuited to inference on chromosome length haplotypes. Therefore, the dominant haplotype likelihood model used for statistical inference is Li and Stephens’ 2003 model (LS) [7] and its various modifications. LS closely approximates the more exact coalescent models but admits implementations with rapid runtime.

Orthogonal to statistical models, another important frontier in genomics is the development of the variation graph [10]. This is a structure which encodes the wide variety of variation found in the population, including many types of variation which cannot be represented by conventional models. Variation graphs are a natural structure to represent reference cohorts of haplotypes since they encode haplotypes in a canonical manner: as node sequences embedded in the graph [8].

In this paper, we present the first statistical model for haplotype modelling with respect to graph-embedded populations. We also describe an efficient algorithm for calculating haplotype likelihoods with respect to large reference panels. The algorithm makes significant use of Novak’s graph positional Burrows-Wheeler transform (gPBWT) [8] index of haplotypes.

## 2 Encoding the full set of human variation

Haplotypes in the Kingman *n*-coalescent and Li-Stephens models are represented as sequences of values at linearly ordered, non-overlapping binary loci [5,6,7]. Some authors model multiallelic loci (for example, single base positions taking on values of (*A, C, T, G, or gap*) [6], but all assume that the entirety of genetic variation can be expressed by values at linearly ordered loci.

However, many types of genetic variation cannot be represented in this manner. Copy number variations, inversions or transpositions of sequence create cyclic paths which cannot be totally ordered. Large population cohorts such as the *1000 Genomes* project data [1] contain simple insertions, deletions and substitution at a sufficient density that these variants overlap or nest into structures not representable by linearly ordered sites. Two examples of this phenomenon from 1000 Genomes data (Phase 3 VCF) for chromosome 22 are pictured below.

**Figure 1:**
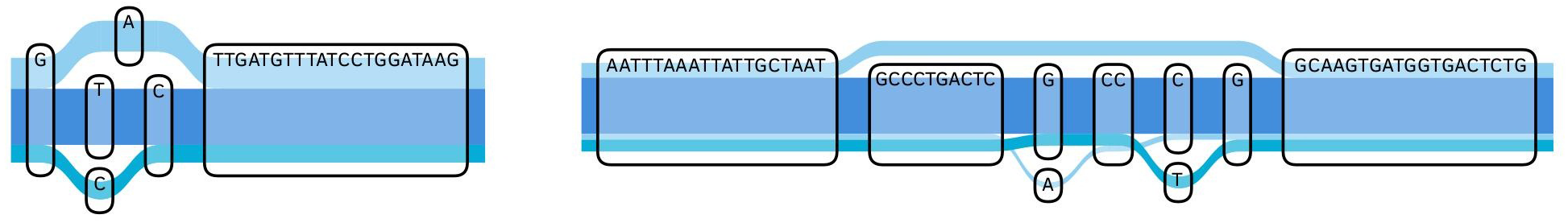
Overlapping, non-linearly orderable loci in a graph of *1000 Genomes* variation data for chromosome 22

In order to represent these more challenging types of variation, we use a *variation graph*. This is a type of *sequence graph*—a mathematical graph in which nodes represent elements of sequence, augmented with 5^′^ and 3^′^ sides, and edges are drawn between sides if the adjacency of sequence is observed in the population cohort [10]. Haplotypes are embedded as paths through oriented nodes in the graph. We are able to represent novel recombinations, deletions, copy number variations, or other structural events by adding paths with new edges to the graph, and novel inserted sequence by paths through new nodes.

## 3 Adapting the recombination component of Li and Stephens to graphs

The Li and Stephens model (LS) can be described by an HMM with a state space consisting of previously observed haplotypes and observations consisting of the haplotypes’ alleles at loci [6,7]. Recombinations correspond to transitions between states and mutations are modelled within the emission probabilities. Since variation graphs encode full nucleic acid sequences rather than lists of sites we extend the model to allow recombinations at base-pair resolution rather than just between loci.

Let *G* denote a variation graph. Let *S*(*G*) be the set of all possible finite paths visiting oriented nodes of *G*. A path *h* in *S*(*G*) encodes a potential *haplotype*. A variation graph posesses an embedded *population reference cohort H* which is a multiset of haplotypes *h* ∈ *S*(*G*). Given a pair (*G*,*H*), we seek the likelihood *P*(*h*|*G*,*H*) that *h* arose from haplotypes in *H* via recombinations.

Recall that every oriented node of *G* is labelled with a nucleic acid sequence. Therefore, every path *h* ∈ *S* corresponds to a nucleic acid sequence *seq(h)* formed by concatenation of its node labels. We represent recombinations between haplotypes by assembling subsequences of these sequences *seq(h)* for *h* ∈ *H*. We call a concatenation of such subsequences a *recombination mosaic*. This is pictured below.

**Figure 2:**
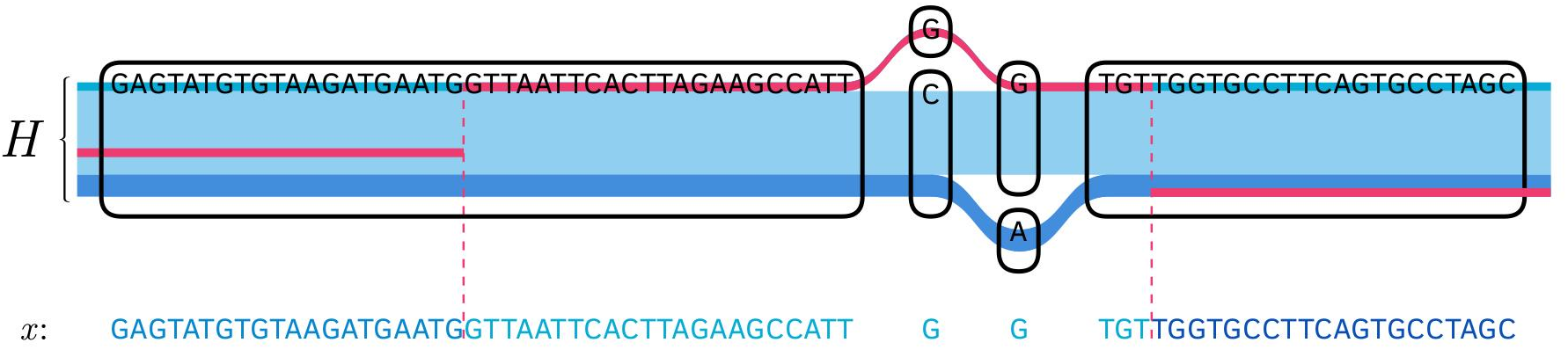
The pink path shows the recombination mosaic *x* superimposed on the embedded haplotypes *H* in our *1000 Genomes project* chr 22 graph; below *x* is mapped onto its nucleic acid sequence.

We can assign a likelihood to a mosaic *x* by analogy with the recombination model from Li and Stephens. Assume that nucleotide in *x* has precisely one successor in each *h* ∈ *H* to which it could recombine. Then, between each base pair, we assign a probability π_*r*_ of recombining, and therefore a probability (1– (|*H*| − 1) π_*r*_) of not recombining. Write π_*c*_ for (1– (|*H*| −1) π_*r*_).

By the same argument underlying the Li and Stephens recombination model, we then we have a probability of a given mosaic having arisen from (*G,H*) through recombinations:

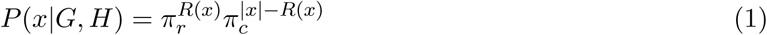

where |*x*| is the length of *x* in base pairs and *R(x)* the number of recombinations in *x*. We will use this to determine the probability *P*(*h*|*G*, *H*) for a given *h* ∈ *S*(*G*), noting that multiple mosaics *x* can correspond to the same node path *h* ∈ *S*(*G*).

Given a haplotype *h* ∈ *S*(*G*), let 
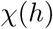
 be the set of all mosaics involving the same path through the graph as *h*. The law of total probability gives

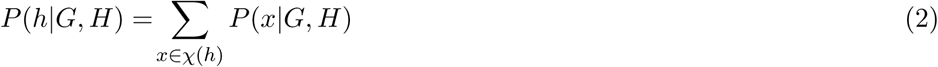

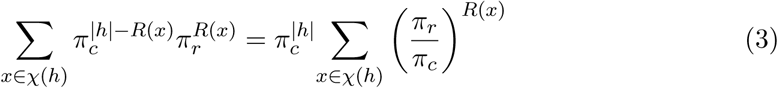

Let 
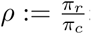
 then *P(h|G,H)* is proportional to a *ρ^R(x)^*-weighted enumeration of 
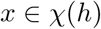

We can extend this model by allowing recombination rate π(*n*) and effective population size |*H*|_*eff*_(*n*) to vary across the genome according to node *n* ∈ *G* in the graph. Varying the effective population size allows the model to remain sensible in regions traversed multiple times by cycle-containing haplotypes. In our basic implementation we will assume that *π*(*n*) is constant and |*H*|_*eff*_(*n*) = |*H*|, however varying these parameters does not add to the computational complexity of the model.

## 4 A linear-time dynamic programming for likelihood calculation

We wish to calculate the sum 
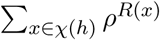
 effciently. (See Eq. 3 above) We will achieve this by traversing the node sequence *h* left-to-right, computing the sum for all prefixes of *h*. Write *h_b_* for the prefix of *h* ending with node *b*.

### Definition 1.

A *subintervals* of a haplotype *h* is a contiguous subpath of *h*. Two subintervals *s*_1_, *s*_2_ of haplotypes *h*_1_, *h*_2_ are *consistent* if *s*_1_ = *s*_2_ as paths, however we distinguish them as separate objects.

### Definition 2.

Given a node *b* of a haplotype *h*, 
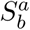
 is the set of subintervals *s*^′^ of *h*^′^ ∈ *H* such that

1. there exists a subinterval *s* of *h* which begins with *a*, ends with *b* and is consistent with *s*^′^
2. there exists no such subinterval which begins with *a* − 1, the node before *a* in *h* (*left-maximality*)

### Deffinition 3.

For a given prefix *h_b_* of *h* and a subinterval *s*^′^ of a haplotype *h*^′^ ∈ *H*, define the subset 
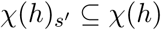
 as the set of all mosaics whose rightmost segment corresponds to *s*^′^.

The following result is key to being able to efficiently enumerate mosaics:

### Claim 1.

*If* 
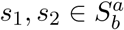 for some a, then there exists a recombination-count preserving bijection between 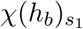 *and* 
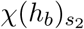.

*Proof*. See supplement.

### Corollary 1.

If we define

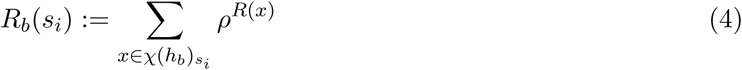

then 
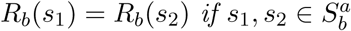 for some a. Call this shared value R_b_(a).

### Definition 4.

*A_b_* is the set of all nodes *a ∈ G* such that 
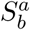
 is nonempty.

Using these results, the likelihood *P*(*h_b_*|*G,H*) of the prefix *h_b_* can be written as

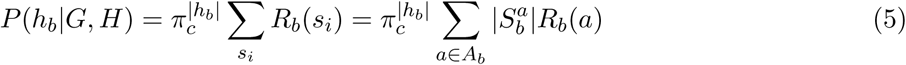

Let *b* – 1 represent the node preceding *b* in *h*; it remains to show that if we know *R*_*b* − 1_(*a*) for all *a* ∈ *A_b – 1_*, we can calculate *R_b_*(*a*) for all *a* ∈ *A_b_* in constant time with respect to |*h*|. This can be recognized by inspection of the following linear transformation:

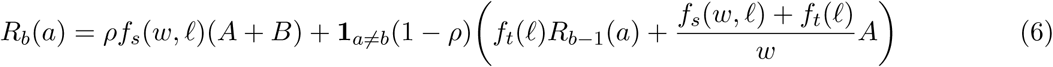

where 
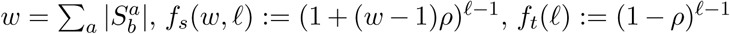
, and *A,B* are the 
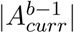
 element sums

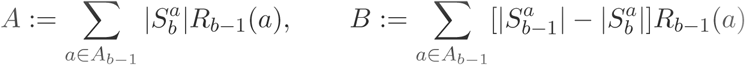

Proof that this computes *R*_*b*_(.) from *R*_*b* – 1_(.) is straightforward but lengthy and therefore deferred to the supplement.

Assuming memoization of the polynomials *f_s_*(*h,ℓ*),*f_t_*(*ℓ*), and knowledge of *w*, *ℓ* and all 
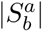
’s, all *R_b_*(*a*)’s can be calculated together in two shared *|A*_*b* – 1_|-element sums (to calculate *A* and *A* + *B*) followed by a single sum per *R_b_*(*a*). Therefore, by computing increasing prefixes *h*_*b*_ of *h*, we can compute *P* (*h*|*G*, *H*) in time complexity which is *𝒪*(*nm*) in *n* = |*h*|, and *m* = *max_b_*|*A_b_*|. The latter quantity is bounded by |*H*| in the worst theoretical case; we will show experimentally that runtime is sublinear in |*H*|.

**Figure 3:**
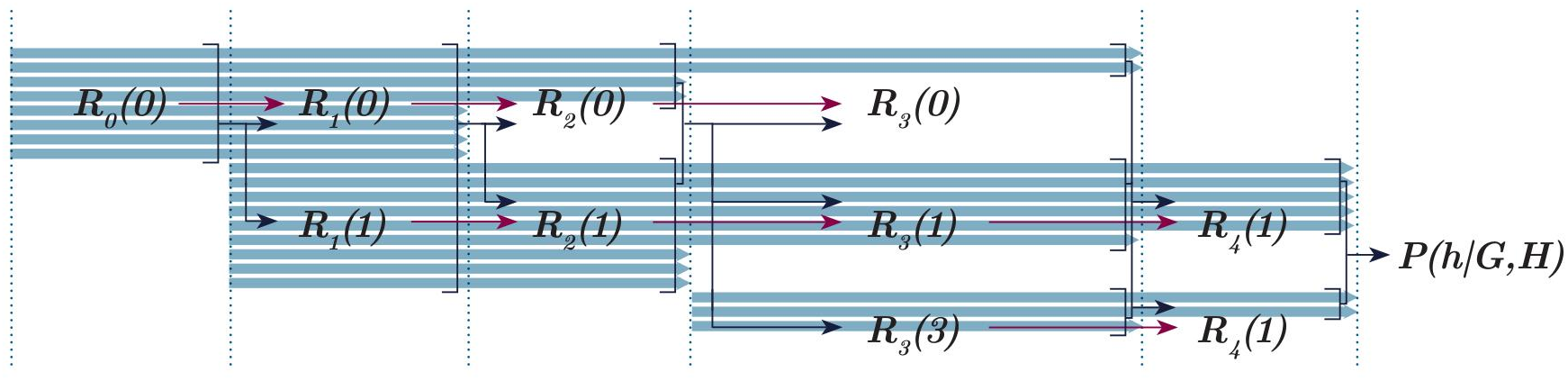
A sketch of the flow of information in the likelihood calculation algorithm described. Blue arrows a represent the *rectangular decomposition*, *R* (.) are prefix likelihoods

## 5 Using the gPBWT to enumerate equivalence classes in linear time

Novak’s graph positional Burrows Wheeler transform index (gPBWT) [8] is a succinct data structure which allows for linear-time subpath search in a variation graph. This is graph analogue of Durbin’s positional Burrows Wheeler transform [3] used in Lunter’s fast implementation of the Viterbi algorithm in the LS model [6]. Like other Burrows-Wheeler transform variants, its search function returns intervals in a path index, and therefore querying the number of subpaths in a graph matching a path is also linear-time in path length.

We demonstrate that we also need only perform a maximum of |*A*_*b* – 1_|+1 gPBWT operations and a corresponding amount of arithmetic to calculate all 
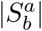
 for a given *b*, provided that we have all 
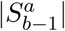

### Definition 5.

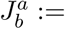
 the number of subpaths in *H* matching *h* between nodes *a* and *b*

This is *𝒪*(*n*) in the gPBWT for *n* = *b* − *a*; in particular, it is *𝒪*(1) given that we have the search interval to compute 
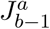
.

### Definition 6.

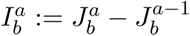

**Claim 2.** 
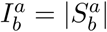

*Proof*. By straightforward manipulation of definitions 2, 5, 6.

It is then evident that |*A*_*b*_ − _1_| + 1 single-node search extension queries^1^ are sufficient to determine 
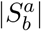
 for all *a* ∈ *A_b_*. Determining these values for all *b* ∈ *h* is therefore also *𝒪*(*nm*) in *n* = |*h*|, and *m* = *max*_*b*_|*A_b_*|

In practice, since 
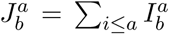
, we can reduce the number of gPBWT queries even further by employing a recursive partitioning of intervals to avoid querying those whose values we can tell are unchanged.

## 6 Modelling mutations

We can assign to two haplotypes *h*, *h*^′^ the probability *P_m_*(*h*|*h*^′^) that *h* arose from *h*^′^ through a mutation event. As in Li and Stephens, we can assume conditional independence properties such that

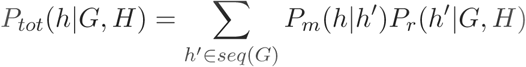

It is reasonable to make the simplifying assumption that *P_mut_*(*h*|*h*^′^) = 0 unless *h*^′^ differs from *h* exclusively at short, non-overlapping substitutions, indels and cycles since more dramatic mutation events are vanishingly rare. This assumption is implicitly contained in the *n*-coalescent and Li and Stephens models by their inability to model more complex mutations.

Detection of all simple sites in the graph traversed by *h* can be achieved in linear time with respect to the length of *h*. The number of such paths remains exponential in the number of simple sites. However, our model allows us to perform branch-and-bound type approaches to exploring these paths. This is possible since we can calculate upper bounds for likelihood knowing either only a prefix, or for interval censored haplotypes where we do not specify variants within encapsulated regions in the middle of the path.

Furthermore, it is evident from our algorithm that if two paths share the same prefix, then we can reuse the calculation over this prefix. If two paths share the same suffix, in general we only need to recompute the 
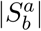
 values for a small number of nodes. This is demonstrated in our evaluation of the time complexity of our methods for haplotypes derived from recombination of previously assessed haplotypes.

## 7 Implementation

To test our methods, we implemented the algorithms described in C++, using elements of the variation graph toolkit *vg* [4] and the gPBWT index implementation in *xg* [8]. This can be found in the “*haplotypes*” branch at http://github.com/yoheirosen/vg. Since no competing graph-based haplotype models exist, we were not able to provide comparative performance data; absolute performance on a single machine is presented instead.

## 8 Results

### 8.1 Runtime for individual haplotype queries

We assessed time complexity of our likelihood algorithm algorithm using the implementation described above. Tests were run on single threads of an Intel Xeon X7560 running at 2.27 GHz.

To assess for time dependence on haplotype length, we measured runtime for queries against a 5008 haplotype graph of human chromosome 22 built from the 1000 *Genomes* Phase 3 VCF on the hg19 assembly created using *vg* and data from the 1000 Genomes project [1]. Starting nodes and haplotypes at these nodes were randomly selected, then walked out to specific lengths. In our graph, 1 million nodes corresponds, on average, to 16.6 million base pairs. Reported runtimes are for performing both the rectangular decomposition and likelihood calculation steps.

**Figure 4:**
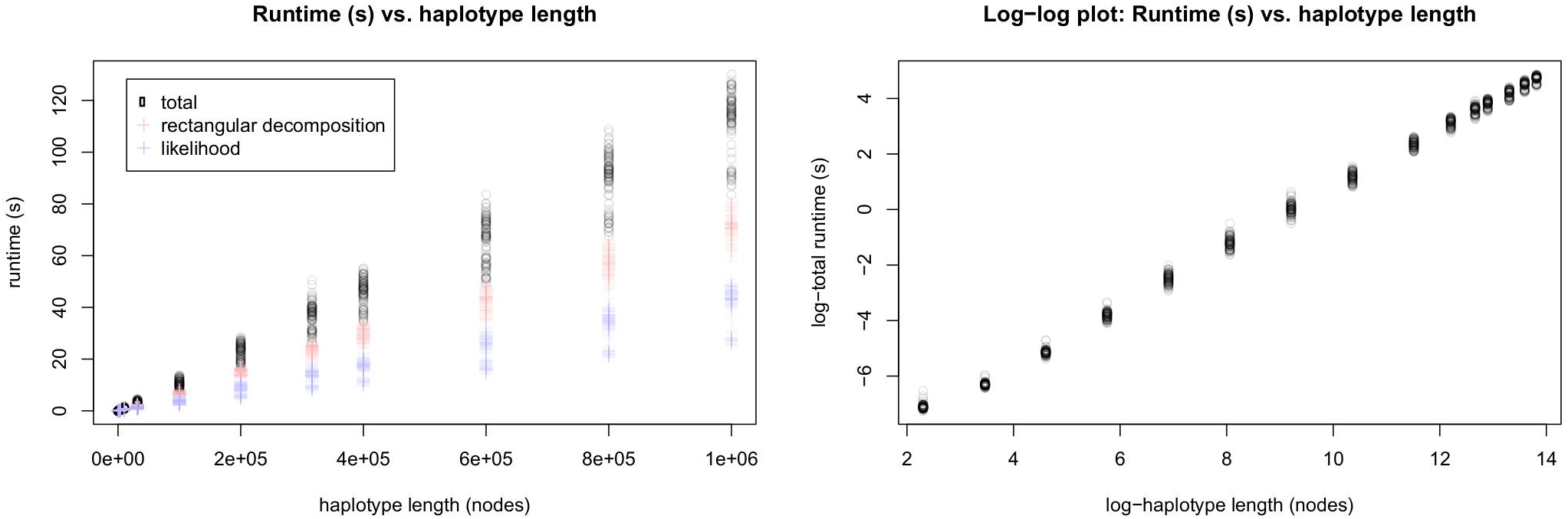
Runtime (s) vs. haplotype length (nodes) for chromosome 22 1000 Genomes data

The observed relationship of runtime to haplotype length is consistent with *𝒪(n)* time complexity with respect to *n* = |*h*|.

We also assessed the effect of reference cohort size on runtime. Random subsets of the *1000 Genomes* data were made using *vcftools* [2] and our graph-building process was repeated. Five replicate subset graphs were made per population size with the exception of the full population graph of 2504 individuals.

**Figure 5:**
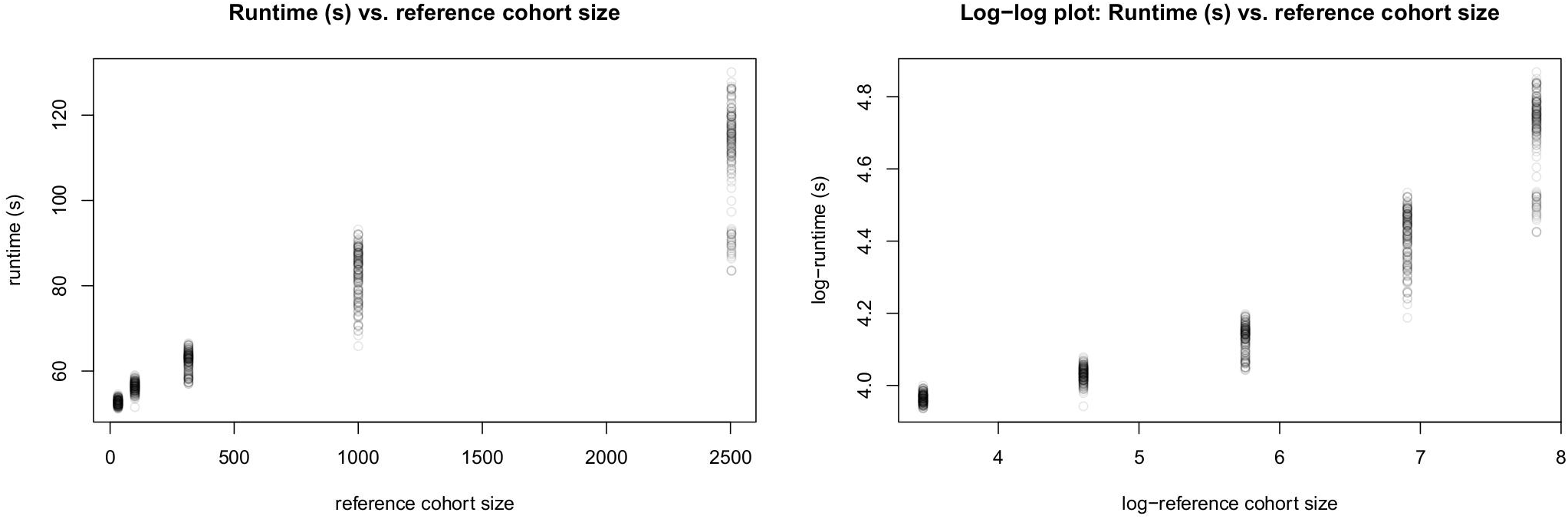
Runtime (s) vs. reference cohort size (diploid individuals) for chromosome 22 1000 Genomes data

We observe an asymptotically sublinear relationship between runtime and reference cohort size.

### 8.2 Time needed to compute the rectangular decomposition of a haplotype formed by a recombination of two previously queried haplotypes

The assessments described above are for computing the likelihood of a single haplotype in isolation. However, haplotypes are generally similar along most of their length. It is straightforward to generate rectangular decompositions for all haplotypes *h* ∈ *H* in the population reference cohort by a branching process, where rectangular decompositions for shared prefixes are calculated only once. This will capture all variants observed in the reference cohort.

Haplotypes not in the reference cohort can then be generated through recombinations between the *h* ∈ *H*. If this produces another haplotype also in *H*, it suffices to recognize this fact. If not, then given that *h* is formed by a recombination of *h*_1_ and *h*_2_, then *h* must contain some sequence of nodes *c* → *j* contained in neither *h*_1_ nor *h*_2_. We only need to recalculate 
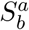
 for *a* ≤ *j* ≤ *b*.

We have implemented methods to recognize these nodes and perform the necessary gPBWT queries to build the rectangular decomposition for *h*. The distribution of time taken (in milliseconds) to generate this new rectangular decomposition for randomly chosen *h*_1_; *h*_2_ and recombination point is shown below.

**Figure 6:**
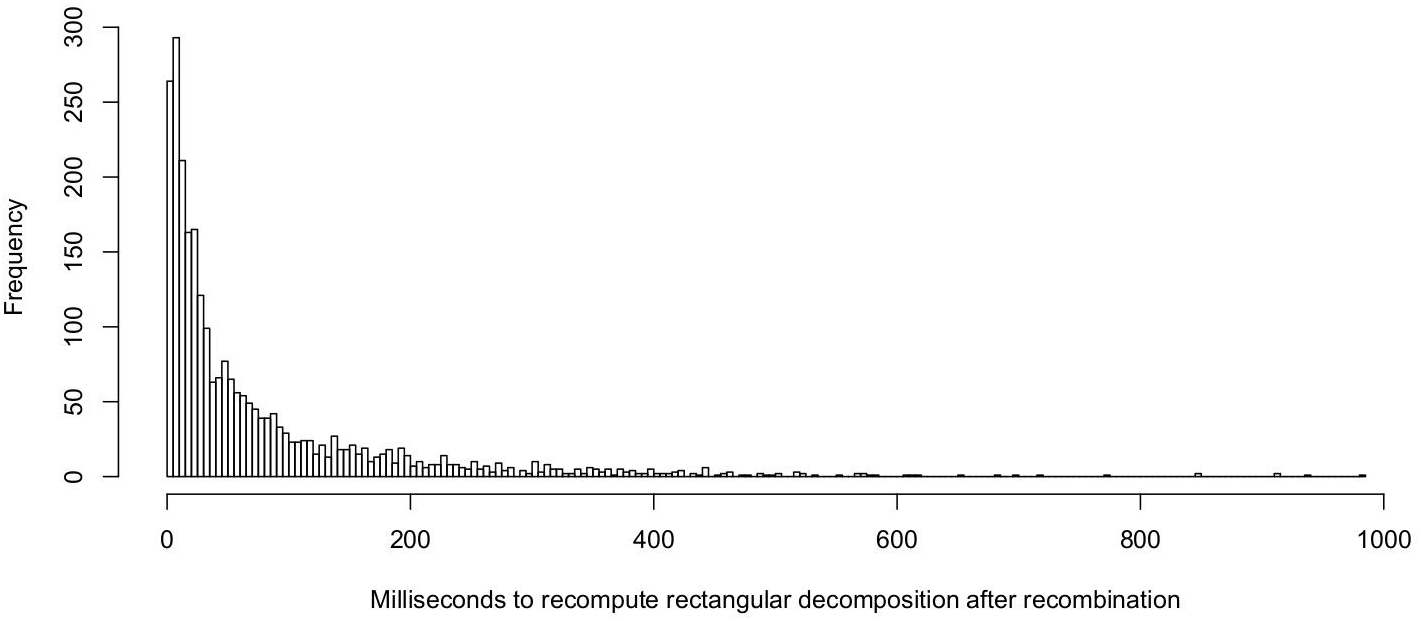
Distribution of times (in milliseconds) required to recompute the rectangular decomposition of a haplotye given that it was formed by recombination of two haplotypes for which rectangular decompositions have been constructed. This graph omits 0.6% of observations which are outliers beyond 1 second of time.

Mean time is 141 ms, median time 34 ms, first quartile time 12 ms and 3rd quartile time 99 ms. To compute a rectangular decomposition from scratch mean time is 71,160 ms, first quartile time 68,690 ms and 3rd quartile time 73,590 ms.

This rapid calculation of rectangular decompositions formed by recombinations of already-queried haplotypes is promising for the feasibility of a mutation model or of sampling the likelihoods of large numbers of haplotypes. Similar methods for the likelihood computation using this rectangular decomposition are a subject of our current research.

### 8.3 Qualitative assessment of the likelihood function’s ability to reflect rare-in-reference features in reads

We used vg to map the 1000 Genomes low coverage read set for individual NA12878 on chromosome 22 against the variation graph described previously. 1476977 reads were mapped. Read likelihoods were computed by treating each read as a short haplotype. These likelihoods were normalized to “relative log-likelihoods” by computing their log-ratio against the maximum theoretical likelihood of a sequence of the same length. An arbitrary value of 10 ^‒9^ was used for π*_recomb_*.

We define a read to contain *n* “novel recombinations” if it is a subsequence of no haplotype in the reference, but it could be made into one using a minimum of n recombination events. We define the prevalence of the rarest variant of a read to be the lowest percentage of haplotypes in the index which pass through any node in the read’s sequence.

We segregated our set of mapped reads according to these features. We make the following qualitative observations, which can be observed in the plots which follow:

1. The likelihood of a read containing a novel recombination is lower than one without any novel recombinations
2. This likelihood decreases with increasing number of novel recombinations
3. The likelihood of a read decreases with decreasing prevalence of its rarest variant

**Figure 7:**
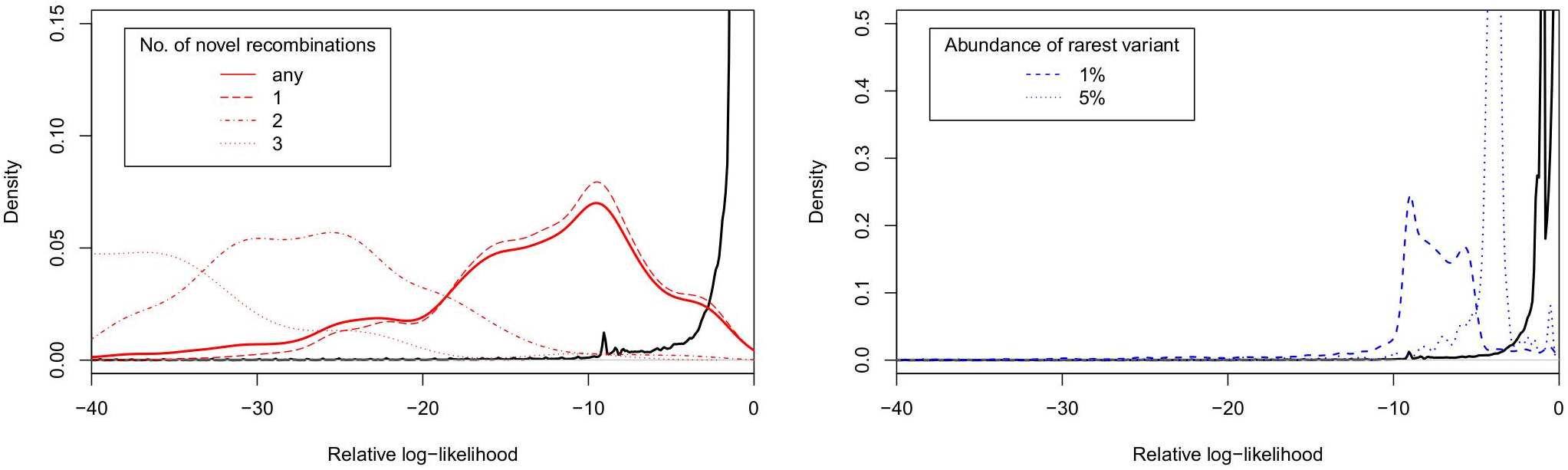
Left: density plot of relative log-likelihood of reads not containing variants below 5% prevalence or novel recombinations (black line) vs reads containing novel recombinations. Right: density plot of relative log-likelihood of reads not containing variants below 5% prevalence or novel recombinations (black line) vs reads containing variants present at under 5% prevalence and under 1% prevalence.

A further comparison of these same mapped reads against reads which were randomly simulated without regard to haplotype structure shows that the majority of mapped reads from NA12878 score are assigned higher relative log-likelihoods than the majority of randomly simulated reads.

**Figure 8:**
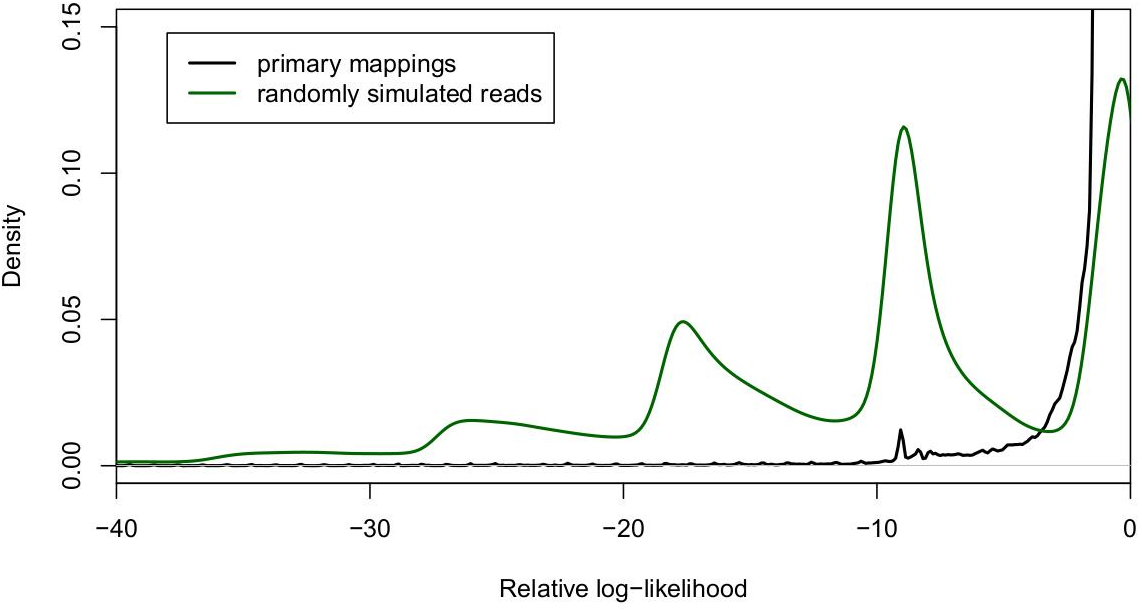
Density plot of relative log-likelihood of mapped reads versus randomly generated simulated haplotypes

## 9 Conclusions

We have introduced a method of describing a haplotype with respect to the sequence it shares with a variation graph-encoded reference cohort. We have extended this into an efficient algorithm for haplotype inference based on Novak’s gPBWT [8]. We applied this method to a full-chromosome graph consisting of 5008 haplotypes from the 1000 Genomes data set to show that this algorithm can efficiently model recombination with respect to both long sequences and large reference cohorts. This is an important proof of concept for translating haplotype inference to the breadth of genetic variant types and structures representable on variation graphs.

Our implementation does not model mutation. This depends on being able to efficiently calculate likelihoods for similar haplotypes. We demonstrate that we can compute rectangular decompositions for haplotypes related by a recombination event in millisecond-range times. We have also devised mathematical methods for recomputing likelihoods of similar haplotypes which take advantage of analogous redundancy properties, however, they have yet to be implemented and tested. However, we anticipate that we will be able to compute likelihoods of large sets of related haplotypes on a time scale which makes modelling mutation feasible.

## 10 Acknowledgements

Y.R. is supported by a Howard Hughes Medical Institute Medical Research Fellowship. This work was also supported by the National Human Genome Research Institute of the National Institutes of Health under Award Number 5U54HG007990 and grants from the W.M. Keck foundation and the Simons Foundation. The content is solely the responsibility of the authors and does not necessarily represent the official views of the National Institutes of Health.

## 11 Appendix A: An *𝒪(n.m)* implementation of the rectangular decomposition construction

Suppose that we wish to find subhaplotypes embedded in the graph which are consistent with a query sequence *h* of nodes. In brief, in the gPBWT, indexing information for haplotypes is stored in such a manner that this can be achieved by calling a function StartSearchAtNode(*Node*) on the first node of *h*, which returns a search interval gPBWTInt of a form analogous to the search interval of a Burrows–Wheeler Transform based index of sequences. This search interval is extended by calling an operation Extend(*gPBWTInt,Node*) to extend this search with each additional node in *h*. Finally, this search interval can be converted into a count of matching subhaplotypes using a function Count(gPBWTInt). It is shown in [8] that StartSearchAtNode, Extend and Count all admit *𝒪*(1) implementations.

It is evident that this search process yields a function CountHaplotypeMatches(*h*) which is *𝒪(n)* in the length |*h*| of *h* in nodes. Let *h*_1_*h*_2_*h*_3_…*h*_|*h*|_ − _1_*h*_|*h*|_ denote the node sequence of *h*. Using CountHaplotypeMatches we can identify the set *A* of nodes in *h* such that either 
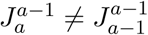
 or 
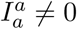
 in *𝒪(n)* independent length-2 subhaplotype count queries:

### Algorithm 1 Identifying *A*, the set of “relevant” nodes

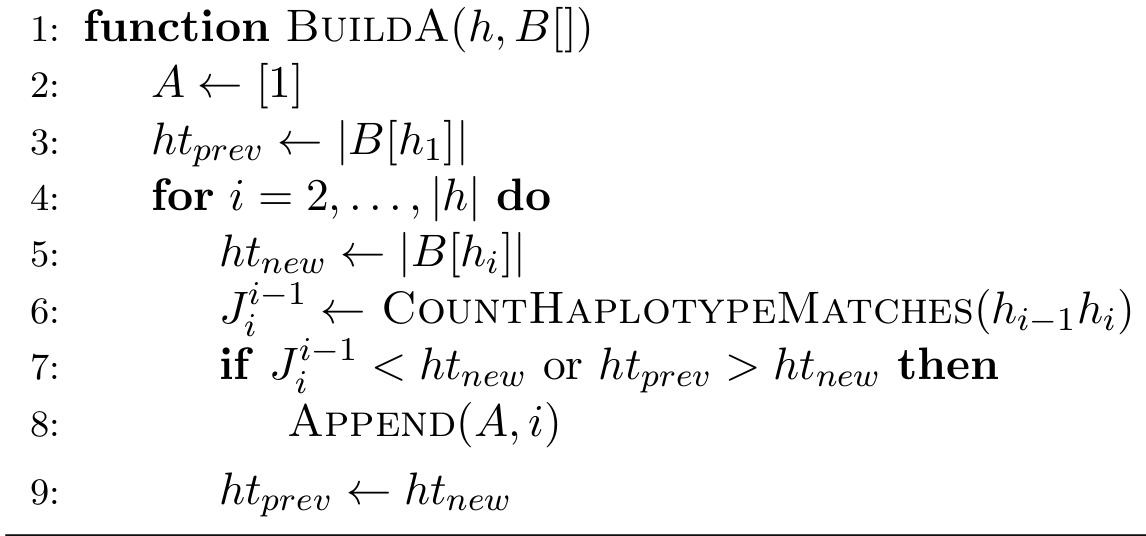

Given that we have constructed *A*, we can determine the rest of the rectangular decomposition and all of the *J*-values according to the following algorithm:

### Algorithm 2 Building the J’s and A_*curr*_’s

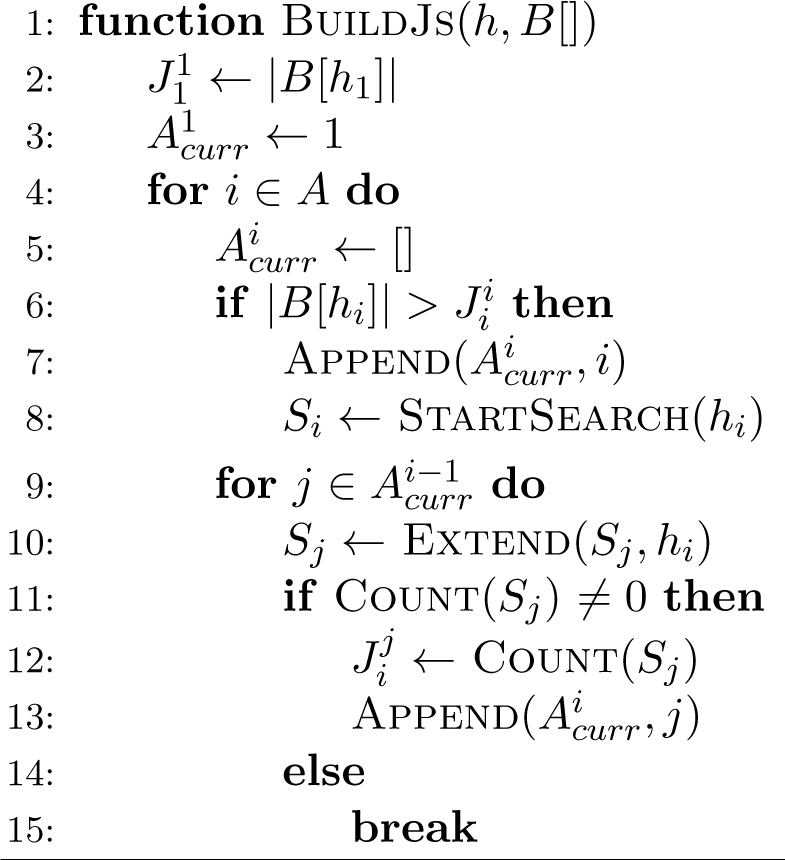

## 12 Appendix B: Arithmetic for derivation of Equation 6

Here we lay out the arithmetic to derive Equation 6, which is used in our iterative computation of likelihood of a haplotype *h* with respect to a population reference cohort *H* embedded in a variation graph *G*. The reasoning is straighforward but involves many subcases which require care.

### 12.1 Notation

#### Definition 7.

A haplotype is a sequence of nodes *n*_1_ → … → *n*_|*h*|_ in a variation graph. The base sequence of a haplotype is the sequence of DNA bases spelled by its node labels. A haplotype subinterval is a contiguous subsequence of a haplotype. A haplotype base sequence subinterval is analogously defined. Denote by |*h*| the length of a haplotype base sequence in base pairs.

#### Definition 8.

Haplotypes *h*, *h*′ are consistent if |*h*| = |*h*′| and *n_i_* = *n_i_′* ∀*i*.

#### Definition 9.

A mosaic of haplotypes *x* consistent with *h* is a vector 〈*x_(i)_*〉 of subintervals of base sequences of haplotypes in *H* whose concatenation is consistent with the base sequence of *h*. The recombination count *R*(*x*) is one less than the number of elements in 〈*x_(i)_*〉. NB: defining these in terms of base sequence rather than node subintervals permits recombination within nodes. Recall Figure 2 from the main text.

#### Definition 10.

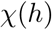
 is the set of all mosaics *x* consistent with *h*. 
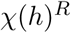
 is the subset with *R*(*x*) = *R*. 
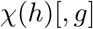
 is the subset whose final subinterval is a subinterval of *g*. 
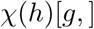
 is that with initial subinterval a subinterval of *g*. 
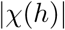
 is the number of elements in 
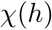
.

### 12.2 Arithmetic shortcuts

#### Lemma 1.

There exists a partition of h into subintervals h_1_, h_2_, …, h_n_ such that if a haplotype g ∈ H has a subinterval consistent with a subinterval of h_i_ then it has a subinterval consistent with all of h_i_.

*Proof.*

It is straightforward to verify that the intervals between successive nodes in the set *A* described in the main text produce such a partition of *h*.

This is important because we will show that it is simple to calculate 
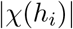
 within any interval with this property.

The following is a more notationally precise statement of Lemma 1 from the main text:

#### Lemma 2.

For any b ∈ A; a ≤ b, given that f and g are members of the same equivalence class 
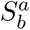
 of haplotypes, the haplotype mosaics 
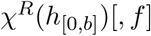
 and 
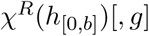
 consistent with the subinterval h_[0,b]_ and ending with subintervals of f and g are in bijective correspondence.

*Proof.*

We assume that *g* ≠ *f* else this is trivial. Consider any mosaic *x* in 
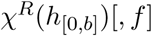
. Given *x* = 〈*x*_1_, *x*_2_,…, *x*_*R*+1_〉, let *j* = max{*i* ∈ 1, …, *R*|*x_i_* is not a subinterval of *g* or *f*}. We will construct a mosaic *y* = 〈*y*_1_; *y*_2_,…, *y*_*R*+1_〉 such that for all *i* ≤ *j*, *y_i_* = *x_i_*, and for all *i* > *j*, *y_i_* is the subinterval of the same length as *x_i_* but derived from the opposite haplotype of the pair *f, g.*

The concatenation *y*_1_*y*_2_,…,*y*_*R*+1_ is consistent with *h*_[0,*b*
]_ since given that both *f*, *g* 
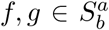
 the first node of *y*_*j*+1_ must be at or after *a*. Therefore clearly 
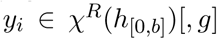
 since its final subinterval corresponds to *g*. The inherent invertibility of this transformation proves that it is a bijection.

**Figure 9:**
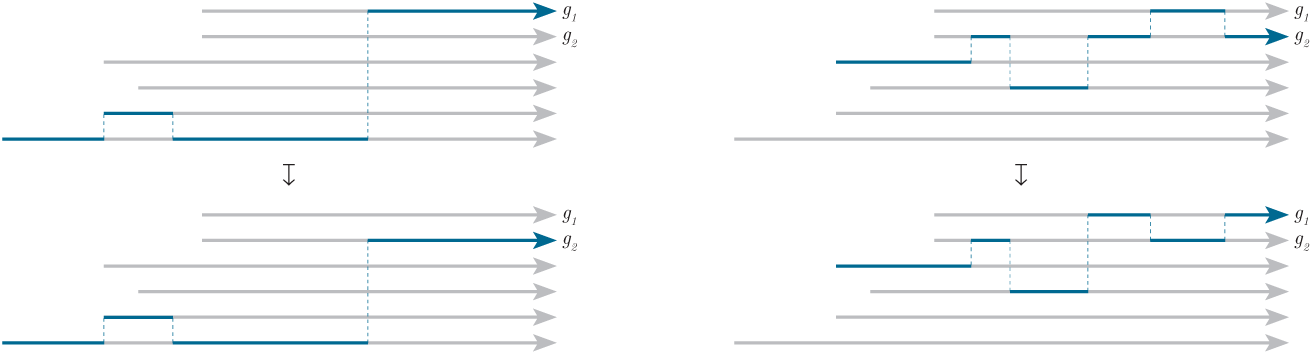
Visual proof of the above lemma by explicit construction of the bijection involved

#### Lemma 3.

Suppose that h_i_ is a subinterval of h such that if a haplotype g ∈ H has a subinterval consistent with a subinterval of h_i_ then it has a subinterval consistent with all of h_i_. Then suppose that f_1_, f_2_, g ∈ H, and all have subintervals consistent with all of h_i_. Then for all R < |h_i_| there is a bijection between 
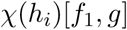
 *and* 
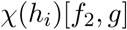

*Proof.*

The proof imitates that of the previous lemma.

### 12.3 The case of a single simple interval *h_i_*

Suppose that *h*_*i*_ is an interval of the form in Lemma 2, *ℓ* base pairs in in length, and has subintervals of *ht* haplotypes of *H* consistent with it. Consider *f*, *g* ∈ *H* such that both have subintervals consistent with *h*_*i*_. Suppose that we wish to calculate, for some *R* < *ℓ*, the number 
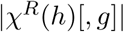
 of mosaics consistent with *h*_*i*
_ having *R* recombinations and ending with haplotype *g*. To calculate 
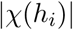
 within an interval of the form above, we need only calculate

1. 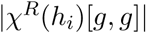
,the number of paths both beginning on and ending on *g* and
2. 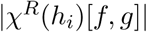
,the number of paths beginning on *f* ≠ *g* and ending on *g*, which, by lemma 4, is the same for all such *f*.

Consider *R* = *ℓ* − 1. It is clear that

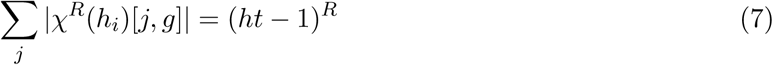

Lemma 4 tells us that all haplotypes *f* ≠ *g* are equivalent for the purposes of enumeration, therefore we write ¬g to denote any arbitrary representative *f* ≠ *g*. There are *ht* − 1 such haplotypes.

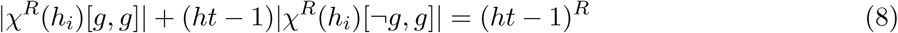

We begin by calculating 
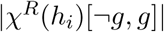
.Consider first *ℓ* = *R* + 1 = 1, for which, given the lack of possible recombinations, 
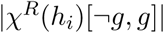
. For *ℓ* = *R* + 1 = 2, any 
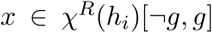
 must at its second node visit a haplotype which is neither *g* nor the ¬g under consideration, therefore 
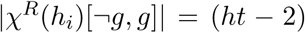
. Suppose now that, for arbitrary *ℓ* = *R*, we know 
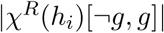
 Then, counting the (*ht* –1) possible haplotypes before finally recombining to *g* shows us that

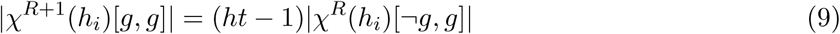

By (8), we know that

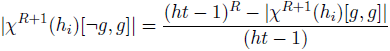

Which by (9) implies

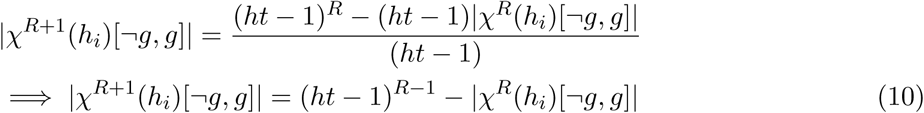

Using (10) as the induction step with base case *ℓ* = *R* + 1 = 2 wend that *∀ℓ* = *R* + 1≥2

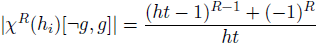

We now relax the restriction that *R* = *ℓ*. For given *R* < *ℓ* each subset of nodes at which recombinations happen will define an additional set of possible recombinations. Counting all possible such subsets

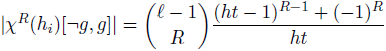

and 
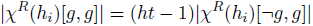

### 12.4 Extending a computation for a prefix by a simple subinterval *h_i_*

To extend our ability to calculate 
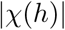
 beyond the single interval *h*_*i*_, suppose we have a partition {*h*_1_, *h*_2_,…, *h_n_*} of *h* into subintervals of the form in Lemma 2. Let *b* ∈ *A* such that b is a node on the boundary of such an interval, let *h*_[0,*b*−1]_ be the prefix of *h* formed by concatenation of the subintervals preceding node *b*, and let *h*_[*b*−1,*b*]_ be the subinterval beginning with node *b*.

Suppose now that we have calculated each 
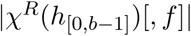
 and now wish to calculate these values up to *b*, the node in *A* succeeding *b* − 1. By Lemma 2, the intervening sequence *h*_[*b* − 1,*b*]_ is of the form for which we have just calculated 
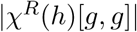
and 
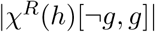
We divide this into cases.

*Case 1*: Suppose that *f* has no subinterval consistent with *h*_[*b,b*+1]_, that is, 
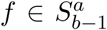
for some *a* but 
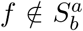
Then any mosaic extending any mosaic in 
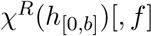
must recombine. Since 
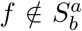
there are 
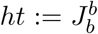
 possible haplotypes to which this recombination at *b* – 1 → *b* may occur. Let *ℓ* be the length (in base pairs) of the interval *b* − 1 to *b*, then ∀*R*^′^< *ℓ(b)* we have previously calculated in (8) that

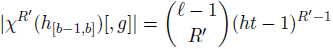

and therefore, where we write 
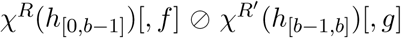
 for the set of mosaics formed by continuing mosaics in 
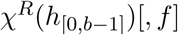
 such that they recombine between *h*_[0,*b* – 1]_ and *h*_[*b* – 1,*b*]_ and end with a subinterval of *g*,

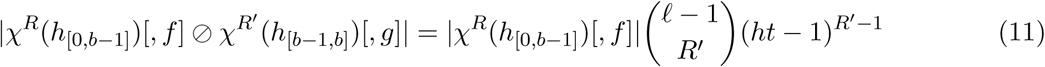

*Case 2*: Suppose now that we know 
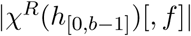
, and 
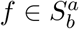
 for some *a*, that is, *f* does have a subinterval consistent with *h*_[*b* – 1, *b*]_

There are two subcases: either there is, or there is not a recombination between the last base in *h*_[0,*b* – 1]_ and the subsequent base at the beginning of *h*_[*b*_ – _1,*b*]_. Suppose that there is not. In this case, where we write 
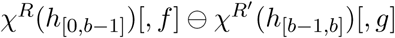
 for the set of mosaics formed by continuing mosaics in 
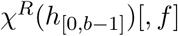
 such that they do not recombine between *h*_[0,*b* – 1]_ and *h*_[*b* – 1,*b*]_ and such that they do end with a subinterval of *g,*

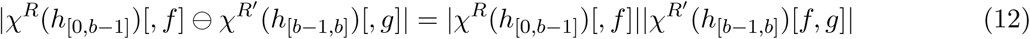

such that if *f* ≠ *g*

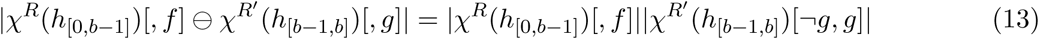

else

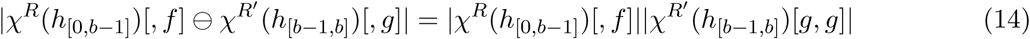

The other subcase is that there is a recombination between the last base in *h*_[0,*b* – 1]_ and the subsequent base at the beginning of *h*_[*b* – 1,*b*]_. In this case if *f* ≠ *g*,

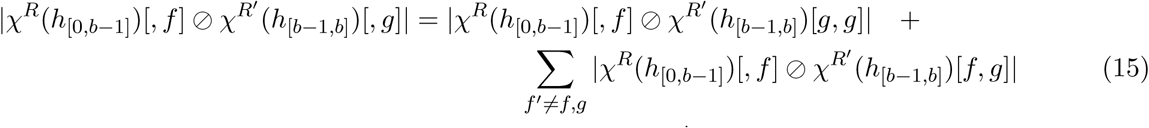

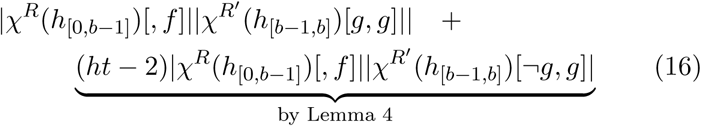

else

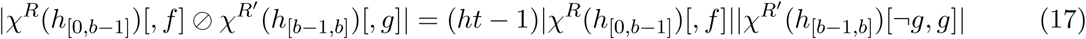

### 12.5 Deriving the Formula for *P*(*h*|*G*,*H*)

Suppose that we have calculated 
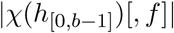
 for all *f* and now wish to calculate 
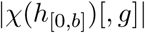
 for some 
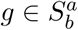
 for some *a≤b*

Note that as defined in the main text, 
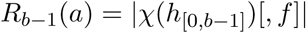
 for the *a* such that 
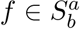
 This means this calculatiuon will in fact give us the formula with which to calculate 
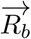
 given 
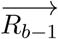
 Let us write *R_b_(f)* for *R_b_(a)* such that 
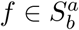

Accounting for all prefixes in 
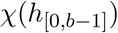
which can produce mosaics in 
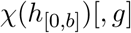then

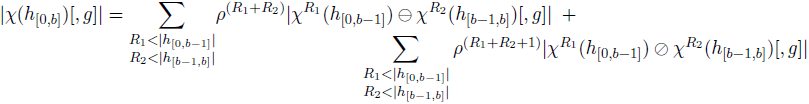

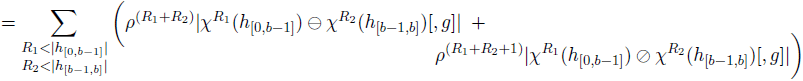

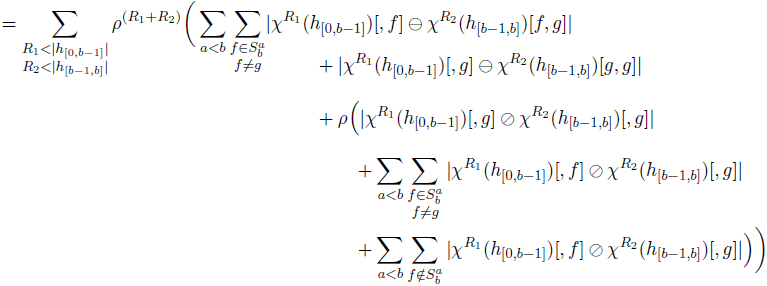

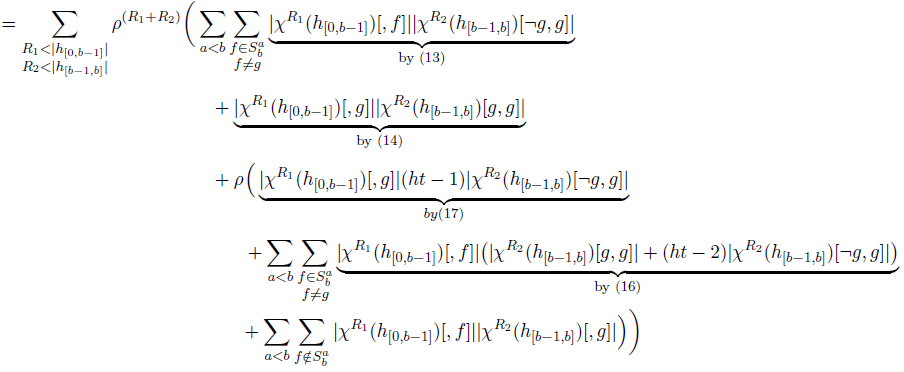

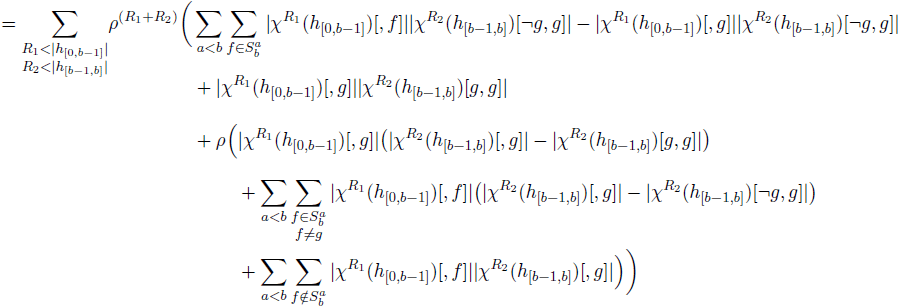

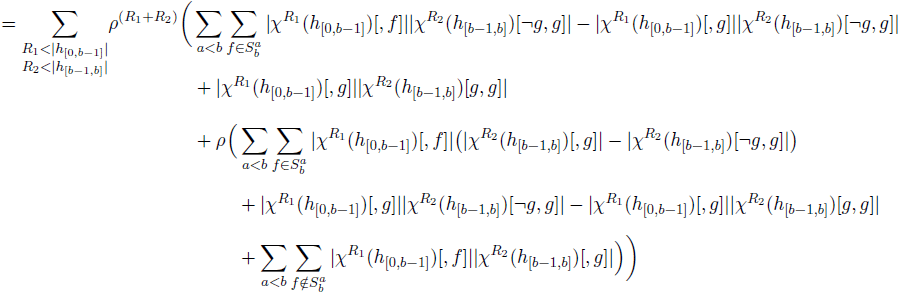

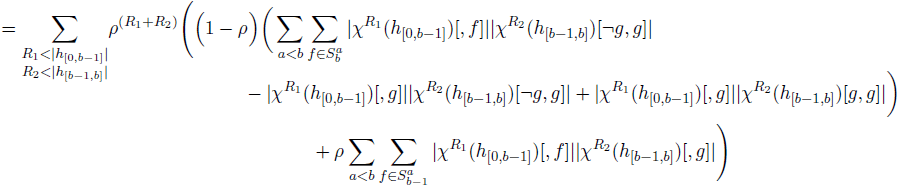

Letting

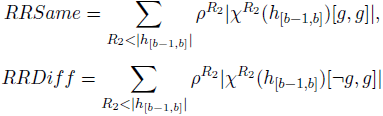

(And we note that *RRSame* and *RRDiff* do not actually depend on choice of *g*)

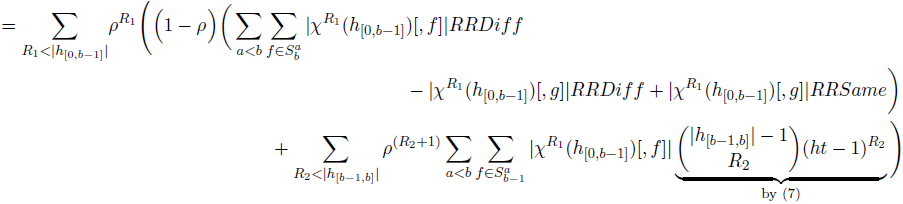

Noting that

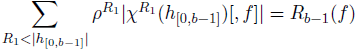

Letting:

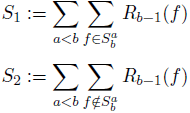

then the above is equal to

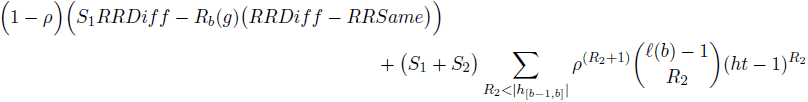

For 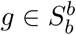 the calculation is similar:

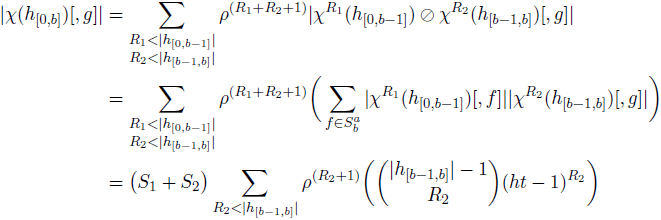

We can simplify the sums above by writing

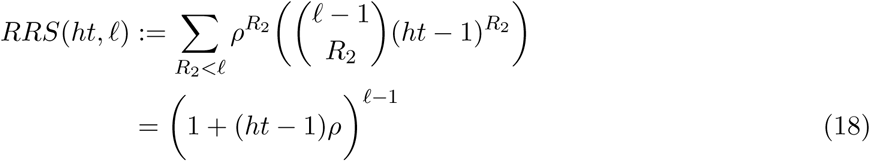

Given a second definition

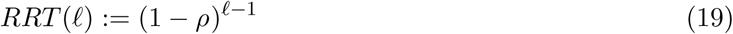

we can actually write

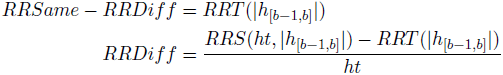

and so finally, we can write our formula for *R_b_(g)* in a compact form as

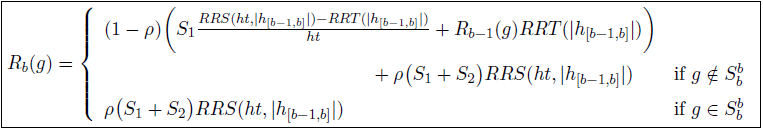

which gives us equation 6 of the main text.

## Footnotes

1 the additional one is to compute 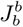

